# Cytokinin synthesis and export from symbiotic root nodules coordinates shoot growth with nitrogen fixation

**DOI:** 10.1101/2022.12.03.518951

**Authors:** Yumeng Chen, Jie Liu, Jieshun Lin, Yuda Purwana Roswanjaya, Marcin Nadzieja, Flavien Buron, Wouter Kohlen, Markus Geisler, Jens Stougaard, Dugald Reid

## Abstract

- Development of symbiotic root nodules is a cytokinin-dependent process that is critical to nitrogen acquisition in legumes. The extent and manner in which root nodules contribute to whole-plant cytokinin and nitrogen supply signalling is unknown.
- Using a combination of genetic, biochemical and physiological approaches, we characterised the role of cytokinin synthesis, export and perception in coordination of symbiotic nodule development and shoot growth in the legume *Lotus japonicus*.
- *LjPup1* encodes a plasma membrane localised cytokinin exporter with isopentenyladenine (iP) and *trans*-Zeatin (tZ) export capacity. *LjPup1* shows a distinct nodule-specific expression pattern with greatest transcript levels detected in mature nodules. Mutants accumulate more isopentenyladenine riboside (iPR) in nodule tissues and demonstrate hallmarks of reduced cytokinin signalling. Despite normal nodule numbers and function, shoot growth is markedly reduced in *Ljpup1* mutants, as well as in mutants impaired in tZ biosynthesis.
- We found symbiotic root nodules contribute to shoot growth via export of active cytokinins. A cytokinin exporter in the purine permease family thus contributes to long-distance cytokinin homeostasis regulating plant development.

## Introduction

Plants constantly adapt their growth in response to nutrient availability. This growth must be coordinated between nutrient acquisition in the roots and the development of aboveground organs. Phytohormones and other mobile signals play critical roles in this process, signalling nutrient availability and demand in systemic circuits (Sakakibara, 2021). In particular, cytokinins act to coordinate nitrate availability, and a number of plant species increase cytokinin signalling following nitrate exposure (Miyawaki et al., 2004; Kamada-Nobusada et al., 2013). Increased cytokinin production in the roots can stimulate shoot growth (Kiba et al., 2013; Osugi et al., 2017; Landrein et al., 2018), including through induction of WUSCHEL (WUS) and increased Shoot Apical Meristem (SAM) activity (reviewed in Cammarata et al., 2019).

Cytokinins are unevenly distributed in plant tissues, with studies in Arabidopsis identifying roots and xylem sap as enriched in *trans*-Zeatin (tZ) and its precursor *trans*-Zeatin riboside (tZR), while shoots and phloem sap have relatively higher isopentenyladenine (iP) type cytokinins (Hirose et al., 2008; Kiba et al., 2013; Osugi et al., 2017). Activation of cytokinin within the shoot meristem is critical to maintaining shoot growth, as demonstrated by the meristem promoting activity of LOG genes that catalyse production of cytokinin bases from precursor ribosides (Kurakawa et al., 2007). This cytokinin activity in the SAM stimulates WUS expression and meristem development and acts both on locally produced cytokinin precursors, as well as systemic cytokinin that signals nutrient availability from the soil (Landrein et al., 2018). The impact of nitrogen availability on cytokinin transport rates can be largely explained by the activity of biosynthesis genes in the root, including IPT3 and CYP735A2 which are regulated by primary nitrate signalling modules in Arabidopsis (Maeda et al., 2018).

Export of cytokinins from the roots occurs through active transport processes, such as through action of xylem loading and phloem unloading by ABCG14 in Arabidopsis (Zhang et al., 2014; Poitout et al., 2018; Zhao et al., 2021). In addition to ABCG family transporters, a number of other gene families transport cytokinins, including Purine Permeases (PUP) (Bürkle et al., 2003; Zürcher et al., 2016) and Equilibrative Nucleoside Transporters (ENT) (Hirose et al., 2005; Sun et al., 2005).

Within the PUP family, PUP1 in rice has been shown to localise to the ER and regulate plant height (Xiao et al., 2020) while OsPUP4 and OsPUP7 are cytokinin regulated transporters and overexpression results in increased grain size (Xiao et al., 2019). In Arabidopsis, AtPUP1 functions as a cytokinin uptake transporter, albeit in a non-specific manner with the protein also capable of adenine transport (Bürkle et al., 2003). Arabidopsis PUP14 regulates cytokinin signalling output via cytokinin uptake at the plasma membrane (Zürcher et al., 2016). PUP family members have also been shown to function in uptake of a number of non-cytokinin purine compounds (Gillissen et al., 2000; Szydlowski et al., 2013). Overall, a number of PUP transporters have been characterised; however, demonstration of transport specificity and direction has not been universally demonstrated.

A major feature of legume root development is the establishment of symbiotic organs, called nodules, to host the nitrogen fixing symbiosis. In the early phases of establishing a nitrogen fixing symbiosis, cytokinin signalling plays critical local roles in the root to induce nodule formation. A number of cytokinin biosynthesis (van Zeijl et al., 2015; Jardinaud et al., 2016; Reid et al., 2017), degradation (Reid et al., 2016), perception (Murray et al., 2007; Held et al., 2014; Boivin et al., 2016; Jardinaud et al., 2016) and signalling components (Tan et al., 2020) have been identified as critical to this process. During these early phases cytokinin export via MtABCG56 is also required for activation of cytokinin signalling (Jarzyniak et al., 2021). The high cytokinin signalling activity achieved by these mechanisms converges on the activation of NIN, a key regulator of nodule development (Liu et al., 2019). Once the nodule is established, cytokinin signalling output as evident by transcript and promoter studies is reduced in *Lotus japonicus* (Held et al., 2014; Reid et al., 2017). Reduced cytokinin biosynthetic gene and receptor transcript levels both appear to contribute to this reduction in signalling output. In legumes, where root development must be balanced with nodule development, symbiotic cytokinin signalling is repressed by nitrate (Lin et al., 2021). How cytokinins contribute to coordinating shoot growth in legumes with varied nitrogen supply from mineral and symbiotic sources therefore remains to be determined.

Here we investigate the role of cytokinin transport by the PUP family in nodules of the legume *Lotus japonicus*. We find *LjPup1* to be highly expressed and to function as a cytokinin exporter from nodules. This cytokinin export from nodules functions to stimulate shoot growth and highlights that cytokinin functions as a nitrogen availability signal to coordinate shoot growth with the symbiotic state in the root.

## Materials and Methods

### Sequence analysis

All *L. japonicus* PUP protein sequences were obtained by BLAST search at lotus.au.dk using both AtPUP14 and LjPUP1 as queries. Sequences were checked for the presence of the purine permease domain (PF16913). Phylogenies were inferred using SHOOT (Emms and Kelly, 2022).

### Cloning and constructs

All primers used for cloning are listed in supplemental table 1. For GUS analyses, approximately 2000bp upstream of the start codon of *LjPUP1* was amplified from *L. japonicus* Gifu genomic DNA. For transport assays a genomic fragment was amplified using primers adding overhangs for goldengate cloning that allowed tagging with GFP at either the N or C terminus. All fragments were cloned into a pDONR vector to create goldengate standard parts (Patron et al., 2015). Constructs for plant transformation were assembled using goldengate cloning (Weber et al., 2011) into a pIV10 vector for *Lotus* hairy roots (Stougaard, 1995) or binary vectors modified to accept GoldenGate clones.

### RNAseq analysis

For statistical analysis of RNAseq, A decoy-aware index was built for Gifu transcripts using default Salmon parameters and reads were quantified using the --validateMappings flag (Salmon version 0.14.1) (Patro et al., 2017). Expression was normalised across all conditions using the R-package DESeq2 version 1.20 (Love et al., 2014) after summarising gene level abundance using the R-package tximport (version 1.8.0). Differentially expressed genes with correction for multiple testing were obtained from this DESeq2 normalised data. Normalised count data and differential expression statistics for all genes is available in supplemental dataset 1.

Gene expression data for figure 1, figure S1 and figure S2 was obtained from publicly available resources at lotus.au.dk.

**Figure 1.**
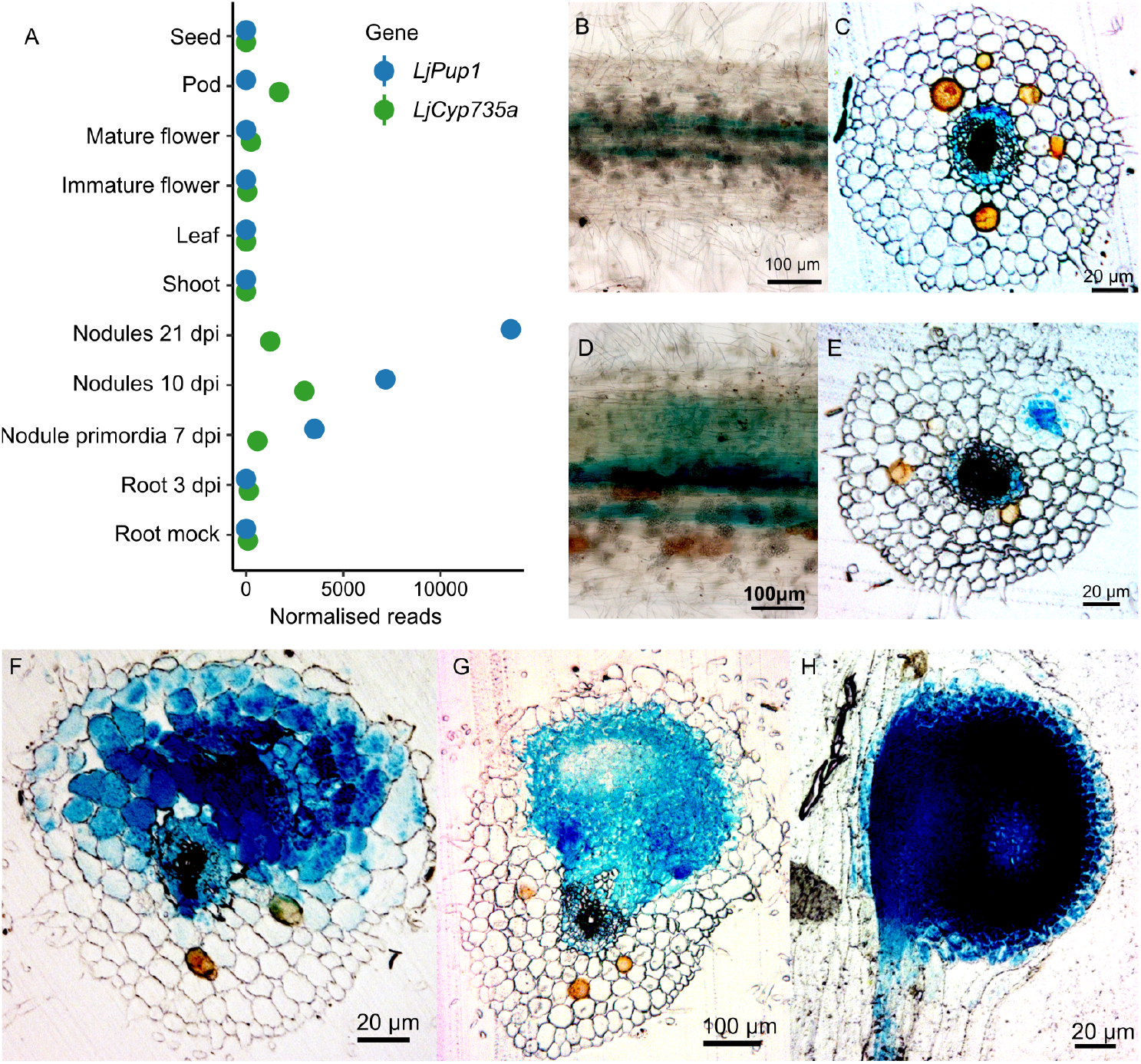
Expression analysis of *LjPup1*. A, RNAseq based expression profile of *LjPup1* and *LjCyp735a* in selected tissues and nodule development series obtained from Lotus Base (lotus.au.dk). B-H *pPup1:GUS* Uninoculated (B-C), 5dpi (D-E), or 20 dpi (F-H) with *M. loti*. B and D are whole mounts, C,E,F,G are transverse sections, and H is a longitudinal section.

### Plant and bacteria growth

*Lotus japonicus* ecotype Gifu was used as wild type (Handberg and Stougaard, 1992), while LORE1 mutants were ordered from LotusBase (https://lotus.au.dk). Homozygotes were isolated for phenotyping as described (Małolepszy et al., 2016). Line numbers and genotyping primers are given in supplemental table 1. *Mesorhizobium loti* R7A was used for nodulation experiments.

Phenotyping on growth plates was conducted by transferring 3-d old seedlings onto filter paper placed on vertical 1.4% agar noble plates containing ¼ B&D media as described (Reid et al., 2016). 3 days after transfer, seedlings were inoculated with rhizobia inoculum OD_600_=0.01. Hairy roots were generated on *L. japonicus* using *Agrobacterium rhizogenes* AR1193 as previously described (Stougaard, 1995).

### *Pup* localisation and Transport assays

For localization experiments, *Pro35S:GFP-PUP1* or *35S Pro35S:PUP1-GFP* was expressed in *N. benthamiana* leaf tissue by *Agrobacterium tumefaciens*-mediated leaf infiltration as described previously (Henrichs et al. 2012). For confocal laser scanning microscopy, a Leica SP5 confocal laser microscope was used and confocal settings were set to record the emission of GFP (excitation 488 nm, emission 500–550 nm) and FM4-64 (excitation 543 nm, emission 580-640 nm).

For protoplast transport assays, protoplasts were prepared from Agrobacterium-transfected *N. benthamiana* leaves and [^14^C]*t*Z and [^3^H]benzoic acid (BA) export was quantified as described previously (Henrichs et al. 2012). In short, tobacco mesophyll protoplasts were prepared 4 days after agrobacterium-mediated transfection of *Pro35S:GFP-PUP1* or *35S Pro35S:PUP1-GFP* or the empty vector control. Protoplast loading was achieved by diffusion and export was determined by separating protoplasts and supernatants by silicon oil centrifugation. Relative export from protoplasts is calculated from exported radioactivity into the supernatant as follows: (radioactivity in the supernatant at time t = x min.) - (radioactivity in the supernatant at time t = 0)) * (100%)/(radioactivity in the supernatant at t = 0 min.); presented are mean values from 4 independent transfections.

For microsomal uptake experiments, *Pro35S:GFP-PUP1* was expressed in *N. benthamiana* leaf tissue by *Agrobacterium tumefaciens*-mediated transfection, and microsomes were prepared and assayed as described previously (Zürcher et al. 2016). In short, ^14^C-labelled *t*Z was added to 300 μg of 25 microsomes in the presence of 5 mM ATP to yield a final concentration of 1 μM [^14^C]*t*Z. For substrate competition assays, unlabelled substrate was included in the transport buffer at a 100-fold excess. After 10 s and 4 min of incubation at 20°C, aliquots of 100 μl were vacuum-filtered and objected to scintillation counting. The indicated relative uptake was calculated as the radioactivity normalized to the first time point (10 s). Means and standard error of means of at least four independent experiments with four technical replicates each are represented.

### GUS staining

GUS staining was performed on transformed hairy roots by vacuum infiltration of GUS buffer (50 mM phosphate buffer, pH 7, 1 mM potassium ferricyanide, 1 mM potassium ferrocyanide, 0.05% (v/v) Triton X-100, and 0.5 mg mL^-1^ 5-bromo-4-chloro-3-indolyl-β-glucuronic acid) followed by incubation at room temperature overnight.

Semi-thin sections of nodules (7 μm) were used for GUS stained tissues. The tissues were incubated in 70% ethanol for 10 days, then were transferred in fixation solution (4% paraformaldehyde and 1% glutaraldehyde) at 4°C for at least 12 hours after 20 minutes of vacuum infiltration. After fixation, the tissues were vacuumed for ∼15 minutes and subsequently dehydrated in an ethanol series of 70%, 80%, 90%, 96% and 96% ethanol for 30 minutes each. For embedding, Technovit 7100 (Heraeus Kulzer) was used according to manufacturer’s instructions; by covering tissue in 25%, 50%, and 75% mix with ethanol under shaking for 1 hour followed by 100% Technovit I overnight. After transferring the tissues into fresh 100% Technovit I solution, the tissues could be embedded with a mix of Technovit I: Hardener II (10:1) until the blocks were hard enough for slicing. Then embedded blocks were sliced into 7 μm layer sections by using a Leica RM2045 microtome.

### Microscopy

Shoots were dissected from Gifu WT and *cyp735a-1* mutants at 14 dpi. To expose the shoot apical meristems, leaves and larger leaf primordia were removed. Images of the SAMs were captured using Zeiss LSM780 confocal microscope by eliciting autofluorescence of the tissue with excitation/emission 405nm/410-562nm parameters. Size of the meristem was defined as its width at widest point and measured using ImageJ.

### Cytokinin measurements

Cytokinin measurements were conducted as previously described (Lin et al., 2021). Approximately 10-20 mg of nodule and shoot tissue was collected from plants 14 days after inoculation, weighed, and snap frozen in liquid nitrogen. Samples were ground and extracted in 1 mL 100% methanol (MeOH) containing stable isotope-labelled internal standards (IS) at 100 nM per compound. Analysis of cytokinins was performed by comparing retention times and mass transitions to the labelled standards, using a Waters Xevo TQs mass spectrometer equipped with an electrospray ionisation source coupled with Acquity UPLC system (Waters, Milford, OH, USA). Chromatographic separations were conducted using an Acquity UPLC BEH C18 column (100 mm, 2.1 mm, 1.7 mm; Waters, Milford, OH, USA) by applying a methanol/water (0.1% formic acid) gradient. Sample concentrations were calculated from a calibration curve for each compound.

### Grafting experiments

Seedlings were grown on ½ strength B5 plates, then 10-day-old seedlings were cut at the hypocotyl and grafted with the aid of plastic tubing fit to the diameter of the stem. After 10 days, grafted plants were planted in Leca in magenta boxes, and 7 days later were inoculated with *M. loti* R7A. Shoot length was analysed 3 weeks post inoculation.

## Results

Cytokinin plays a critical role in initiation and development of symbiotic root nodules in legumes (Murray et al., 2007; Tirichine et al., 2007; Reid et al., 2017). To investigate the role of cytokinin transport in the nodule, we investigated the Purine Permease (PUP) family, several members of which have been shown to transport cytokinin (Bürkle et al., 2003; Zürcher et al., 2016). In *Lotus japonicus*, the PUP family comprises 12 members, with 3 showing significantly elevated transcript levels in root nodules relative to uninoculated roots in the RNAseq-based expression atlas (Fig S1; Kamal et al., 2020). Of these, *Pup1* shows the most dramatic increase in transcript levels, with no transcripts detected in uninoculated root samples, increasing to in excess of 10 000 in mature nitrogen-fixing nodules 21 days post inoculation (dpi) with *Mesorhizobium loti* (Fig 1A). The 10 and 21 dpi nodule samples also exhibit elevated expression of *LjCyp735a*, which catalyses the production of *trans*-Zeatin (tZ; Fig 1A). *Pup4* and *Pup5* also showed elevated transcript levels in the nodule relative to root tissue (Fig S1), although the normalised transcript levels are approximately an order of magnitude lower than *Pup1*. Expression data from mutants impaired in nitrogen fixation (*sst1* and *sen1*) indicate that the onset of nitrogen fixation is not required for the induction of *Pup1* (Fig S2).

A phylogeny for *PUP1* was inferred using SHOOT (Figure S3; Emms and Kelly, 2022), which indicated that LjPUP1 is in an orthogroup containing 6 Arabidopsis proteins: PUP6, PUP7, PUP8, PUP10, PUP21 and PUP22, which are tandemly duplicated in a region of approximately 12.5 kb on Arabidopsis chromosome 4.

To determine the tissue specificity of *Pup1* expression, we cloned a 2 kb promoter region with a GUS reporter and analysed hairy root plants inoculated with rhizobia in a time series. In uninoculated roots, GUS staining was evident in the root vasculature, albeit at a low level (Fig 1B-C). From 5 dpi, staining became evident in the root cortex, and sections revealed this was associated with cell divisions of newly forming nodule primordia (Fig 1D-E). As cell divisions progressed in emerging nodules, staining intensified and was evident throughout the nodule, and in the root vasculature (Fig 1F-G). Mature nodules had very intense staining throughout, and in the adjacent root vasculature (Fig 1H).

To verify the transport activity of *PUP1*, we first sought to identify the protein’s subcellular location. We transfected *Nicotiana benthamiana* leaf tissue by Agrobacterium-mediated leaf infiltration using an N-terminal GFP fusion of PUP1 (GFP-PUP1). Confocal analysis revealed a plasma membrane specific localisation pattern of PUP1 in epidermal cells, which overlayed widely with the plasma membrane marker, FM4-64 (Fig 2A).

**Figure 2.**
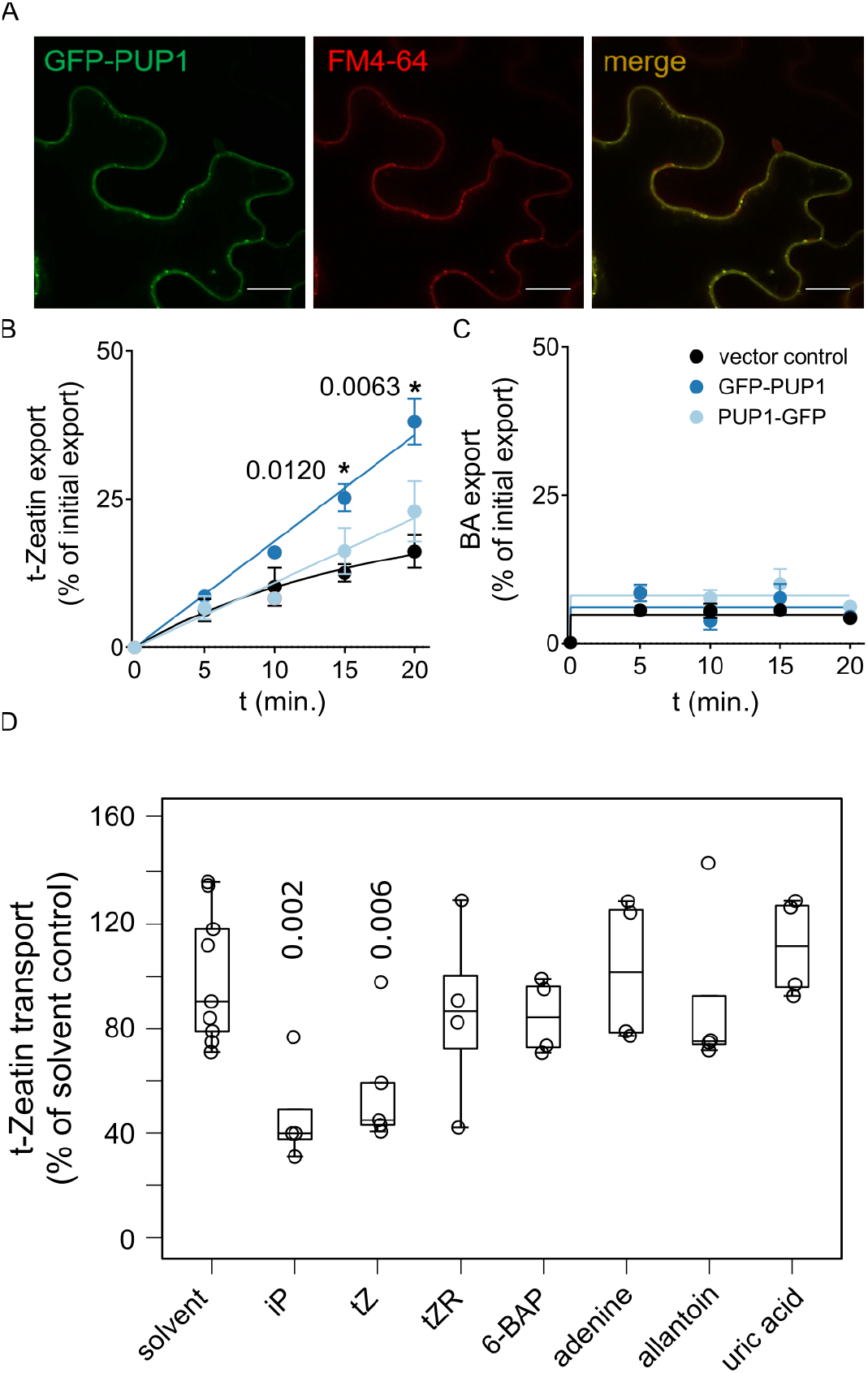
Localisation and transport characteristics of *LjPUP1*. A, Localisation of GFP-PUP1 in *N. benthamiana* leaf epidermal cells. FM4-64 was used as a PM marker; Scale bars, 10 μm. B, Export of [^14^C]*trans*-zeathin (*t*Z) from protoplasts prepared from *N. benthamiana* transfected with GFP-PUP1, PUP1-GFP or the vector control. C, No significant efflux of diffusion control [^14^C]benzoic acid (BA), simultaneously loaded with [^3^H]*t*Z, was observed D, t-Zeatin competition assay using GFP-PUP1 microsomes and indicated competitors at a 100-fold access. Significant differences of means to vector control (**B**) or solvent control (D) are indicated by *P* values and were determined by ordinary One-way ANOVA. *t*Z, *trans*-zeatin; iP, isopentenyladenine; *t*ZR, *trans*-zeatin riboside; 6-BAP, 6-benzylaminopurine

In order to demonstrate cytokinin transport activity for PUP1, we prepared mesophyll protoplasts from tobacco (*Nicotiana benthamiana*) leaves transfected with N-(GFP-PUP1). and C-terminally GFP-tagged PUP1 (PUP1-GFP) or the empty vector control. Expression of both PUP1 constructs resulted in enhanced export of radiolabelled tZ from prepared protoplasts in comparison to the vector control, which however, was only significant for GFP-PUP1 (Fig 2b). A valid explanation might be that C-terminal GFP fusion might lead to inactivation of PUP1 transport activity as described for many transport proteins (REF). This export is specific for the cytokinin, tZ, as the diffusion control benzoic acid (BA) did not show any significant difference to the vector control (Fig 2c). To further investigate the substrate specificity of GFP-PUP1, we tested different cytokinines and adenine derivates (?) in microsomal transport assays upon their potential to compete for transport of radiolabelled tZ. Interestingly, we found that GFP-PUP1 revealed in comparison to other PIPs and ABCG-type cytokinin transporters (REF) a remarkable high substrate specificity with only tZ and iP being able to inhibit significantly tZ import into microsomal fractions. In summary, or data qualify PUP1 as a highly-specific PM exporter of cytokinins.

Having established *Pup1* as a nodule-expressed exporter of cytokinin, we next sought to evaluate the nodulation phenotypes of *pup1* mutants. We obtained 2 LORE1 lines with exonic insertions in the *Pup1* coding sequence (LORE1 IDs *pup1-1* 30059797; *pup1-2* 30066380). Given the tZ export capacity of *Pup1*, we also evaluated mutants in the cytokinin hydroxylase *cyp735a*, which catalyses the formation of tZ from iP-type cytokinins (LORE1 IDs *cyp735a-1* 30079927; *cyp735a-2* 30161349). Nodulation assays indicated that the *pup1-1* allele had a minor, but significant reduction in nodulation, while *pup1-2, cyp735a-1* and *cyp735a-2* showed no nodulation defects relative to wild-type plants (Fig 3A). Given that both *pup1-1* and *pup1-2* contain exonic insertions which are predicted to disrupt gene function, we therefore concluded that the nodulation defect in *pup1-1* is unlikely to result from the mutation in the *Pup1* gene, and more likely reflects one of the 8 additional insertions present in this line. We did not detect any difference in the nitrogen fixation activity of *pup1* or *cyp735a* mutants as determined by acetylene reduction assay (Fig 3B).

**Figure 3.**
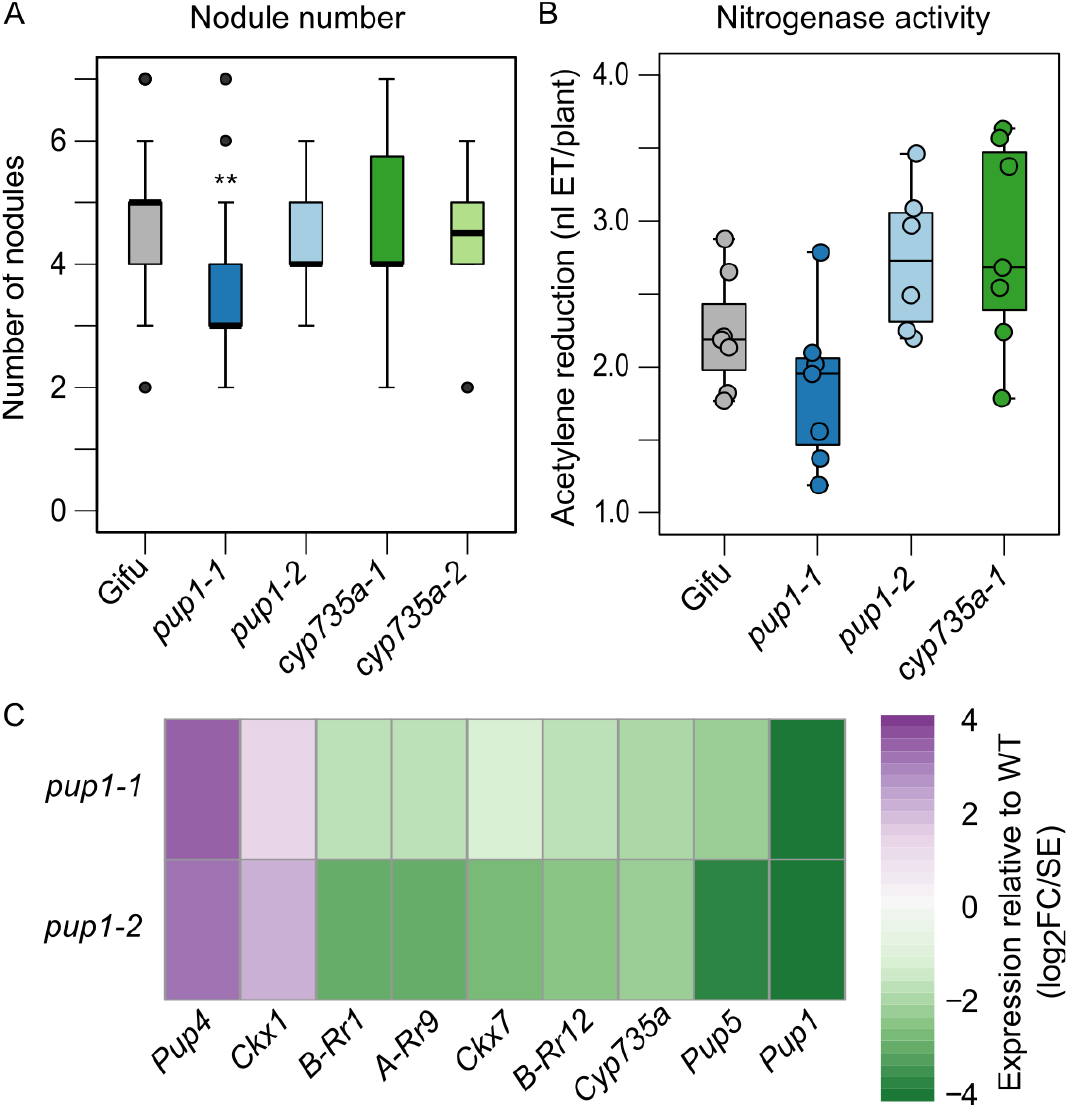
Nodule phenotypes of *pup1* and *cyp725a* mutants. A, Nodule number on plates 21 dpi. B, Nitrogenase activity determined by acetylene reduction assay. C, RNAseq of cytokinin related genes which were differentially regulated in at least 1 *pup* mutant allele. Values shown are the Wald statistic (log_2_ fold-change divided by its standard error) of mutant compared to wild-type nodules. *n*= 14-70 in nodulation assay and 6-7 in acetylene reduction.

To determine the impact of *pup1* gene disruption on cytokinin signalling in the nodule, we conducted RNAseq on nodules harvested at 14 dpi (Fig 3C). This revealed that *pup1* mutant nodules showed a significant compensatory upregulation of *Pup4* and a downregulation of *Pup5* (Fig 3C). We examined the expression of 47 genes associated with cytokinin biosynthesis; breakdown or signalling, and found 5 showed a significant reduction and one an increase in transcript levels when comparing the wild-type with both *pup1* alleles (Fig 3C). The impact in each case was relatively minor.

The altered expression of several cytokinin related genes in *pup* mutant nodules indicates that cytokinin homeostasis may be disturbed in these nodules. We therefore generated transgenic hairy roots of *pup* mutants with a construct carrying the TCSn:GUS reporter as a synthetic readout of cytokinin signalling activity. Analysis of stained nodules revealed a qualitative difference in GUS activity in *pup1* nodules relative to wild-type, with markedly less staining evident (Fig S4 A-H). Sectioned roots showed that this reduction in GUS staining was most evident in the outer cell layers of the nodule (Fig S4c,f,i). We further confirmed the utility of the TCSn:GUS reporter for analysing such qualitative differences by generating transformed roots on the previously characterised *ckx3* mutant, that exhibits elevated cytokinin levels in *L. japonicus* roots and nodules (Reid et al., 2016). This analysis showed that *ckx3* mutants display the opposite impact on TCS activity relative to *pup* mutants, with much more intense GUS staining throughout the nodule (Fig S4j-l).

During our nodulation analysis, we observed that both *pup1* and *cyp735a* mutants tend to exhibit reduced shoot growth relative to wild-type, despite their apparently normal nodulation and nitrogen fixation (Fig 4A). Quantification of shoot length revealed that *cyp735a* mutants are most severely impacted in shoot length, with *pup1* mutants showing an intermediate reduction relative to wild-type (Fig 4B). We performed microscopy on the Shoot Apical Meristem (SAM) to determine if the reduced shoot length was evident at the meristem, but did not identify a difference in SAM size (Fig S5).

**Figure 4.**
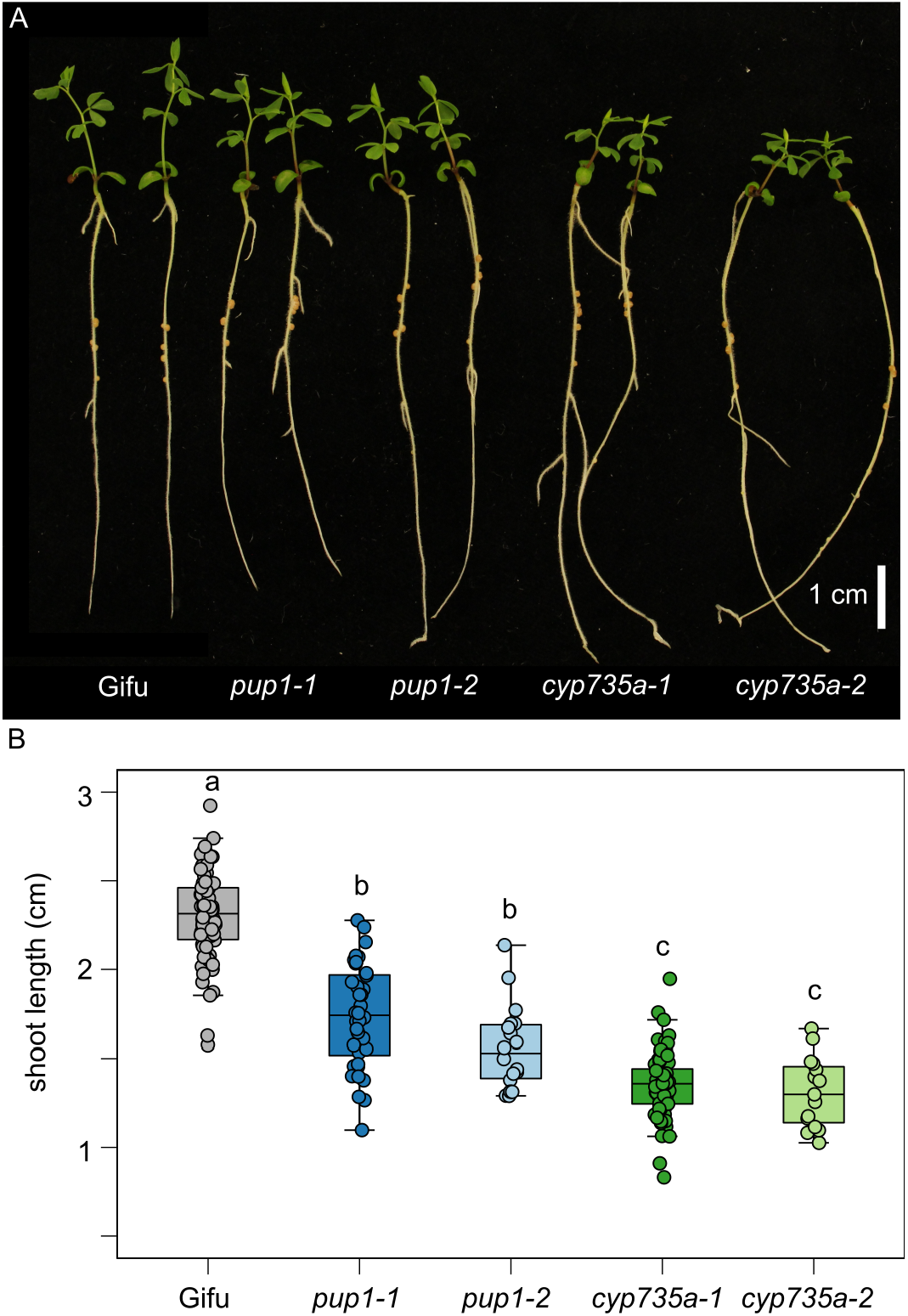
Shoot phenotypes of *pup1* and *cyp725a* mutants. Shoot growth phenotypes of Wild-type, *pup1*, and *cyp735a* mutants grown on nodulation plates in the absence of nitrate (A). Quantification of shoot length conducted from image data for the indicated genotypes (B). Letters indicate significant differences determined using Tukey post-hoc analysis. *n* = 15-70 for shoot length analysis.

To identify the impact of *pup* and *cyp735a* disruption on the nodule and shoot cytokinin levels, we analysed the cytokinin content in the two tissues in the mutants (Fig 5A-B). Analysis of the nodule showed that *pup1* mutants did not show significant differences to wild type in levels of iP, tZ or tZR. On the other hand, levels of iPR were significantly higher in *pup1-1* and *pup1-2* (Fig 5A). Nodules on *cyp735a* mutants showed significantly elevated iP and iPR and reduced tZ and tZR levels (Fig 5A). In the shoot tissue, we did not detect differences in cytokinin in *pup1* mutants, while *cyp735a* mutants showed consistently elevated iP content (Fig 5B). There was a tendency for reduced tZ content in *cyp735a* shoots, with only the *cyp735a-1* allele being significant (Fig 5B).

**Figure 5.**
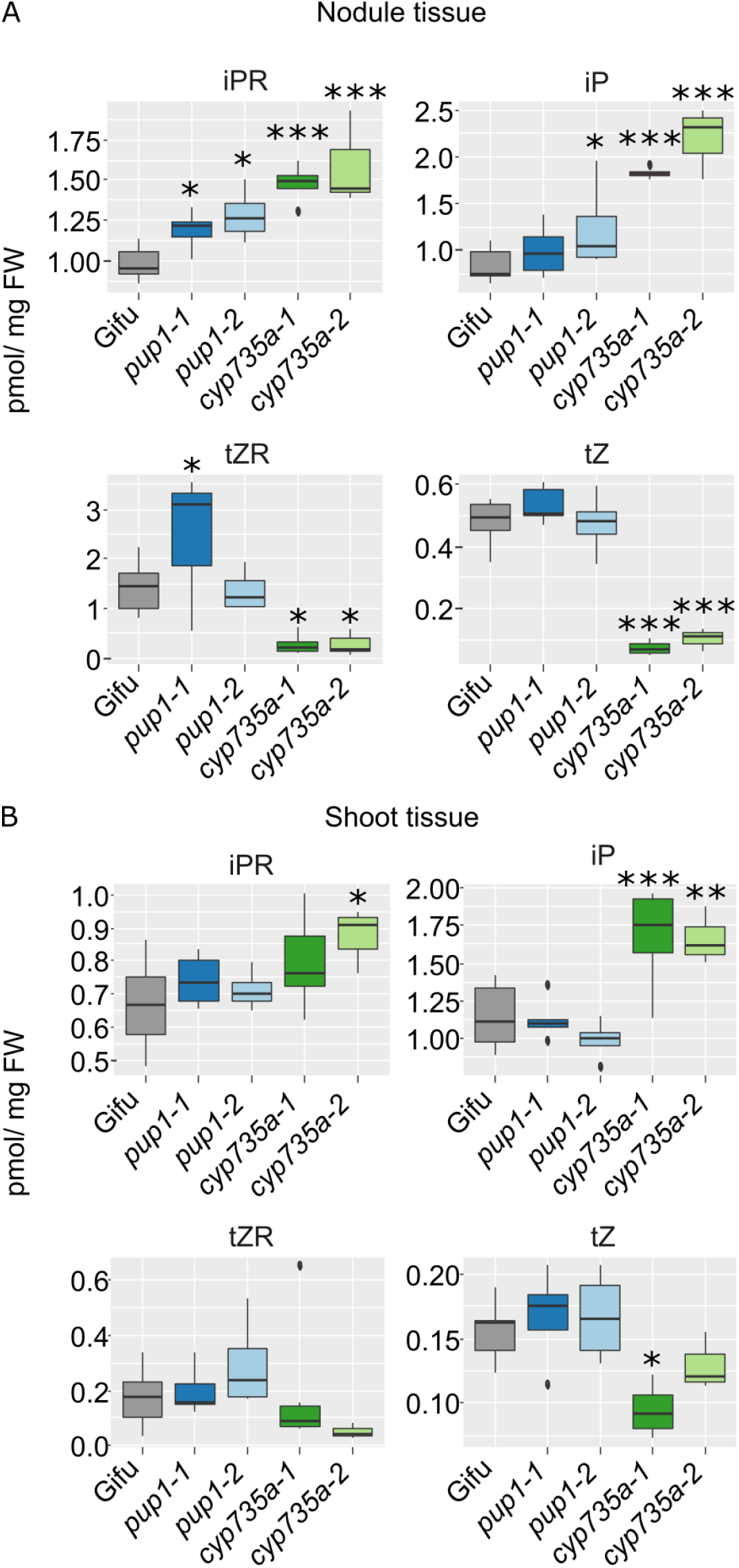
Cytokinin content in nodule and shoot tissues. Nodules (A) and shoots (B) were collected 14 dpi with *M. loti* and indicated cytokinin species measured. Statistical significance determined by planned contrasts against the Gifu control are indicated by *=0.05, **=0.01, ***=0.001. *n* = 4-6

Given the impact on shoot elongation of the cytokinin biosynthesis and transport mutants, we aimed to establish which cytokinin receptors may regulate the shoot phenotypes. In line with previous reports, *Lhk1* transcripts are the most abundant cytokinin receptor in RNAseq data from root tissues (Fig 6A; Held et al., 2014). On the other hand, we found *Lhk2* and *Lhk3* are the most highly abundant transcripts in shoot tissues (Fig 6A). Consistent with previous reports, and the root preferential expression of *Lhk1*, nodulation was significantly reduced in the *lhk1* mutant (Fig 6B). By contrast, the *lhk2 lhk3* double mutant did not display a significant nodulation defect (Fig 6B). Quantification of shoot phenotypes showed *lhk2 lhk3* mutants display severe reductions in length, with *lhk1* having a significant, but intermediate, reduction relative to wild-type and the double mutant (Fig 6C-D).

**Figure 6.**
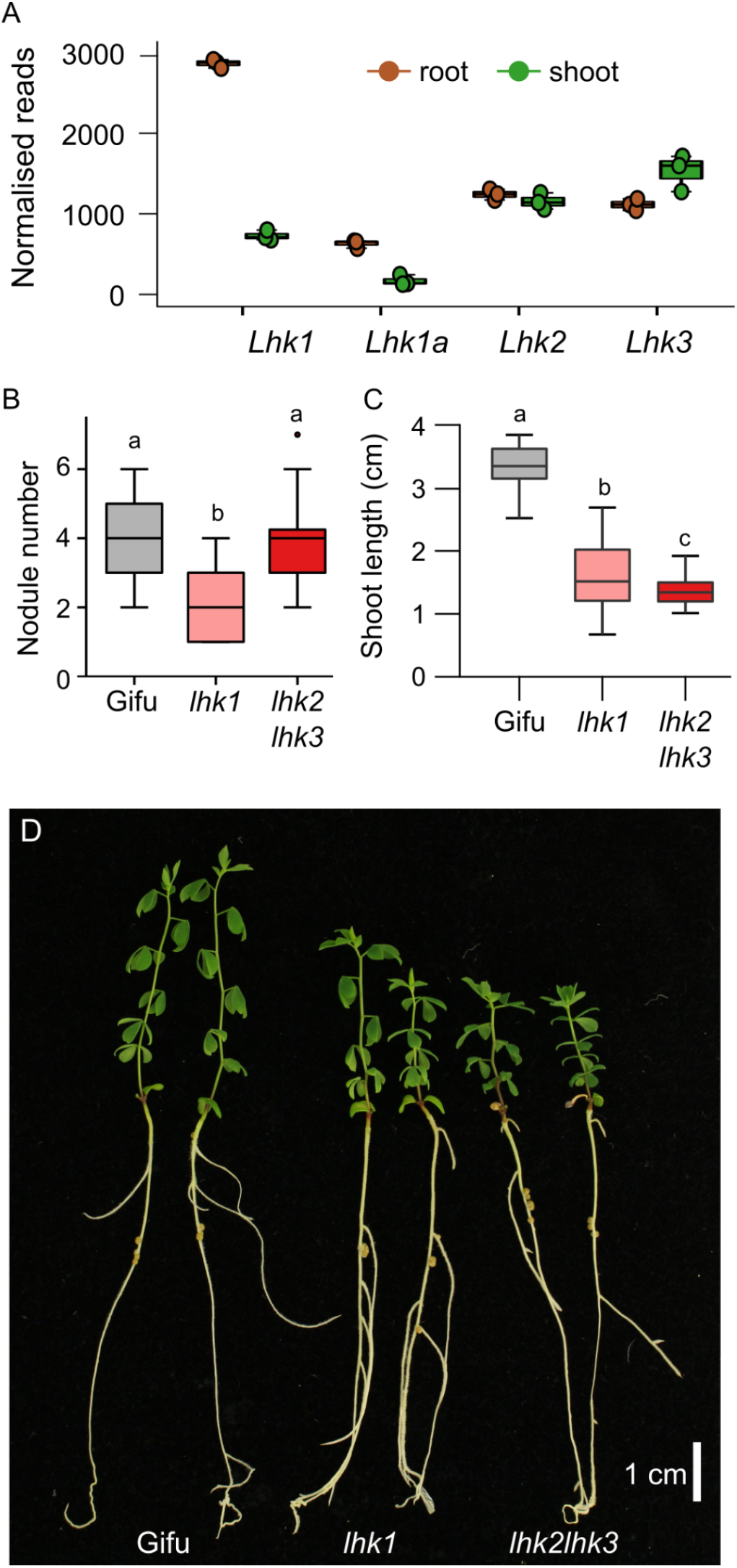
Analysis of root, shoot and nodulation phenotypes of *L. japonicus* cytokinin receptor mutants. A. Normalised RNAseq transcript levels in roots and shoots for *Lhk1, Lhk1a, Lhk2* and *Lhk3* obtained from the RNAseq expression atlas at lotusBase (lotus.au.dk). B. Nodule number on plates 21 dpi with *M. loti* for the indicated genotypes. C. Quantification of shoot length conducted from image data for the indicated genotypes. D. Representative phenotypes for each of the quantified genotypes. Letters indicate significant differences determined using Tukey post-hoc analysis. *n* = 37-40 for nodule and shoot length analysis.

The reduction in shoot growth, coupled with the strictly nodule expression of *Pup1*, indicates that long-distance stimulation of growth may result from nodule-exported cytokinin. To confirm this hypothesis, we quantified shoot growth in grafting experiments with two mutants that show reduced shoot growth despite normal nodulation phenotypes (*cyp735a* and *lhk2 lhk3*, Fig 7). The grafting experiments showed that *Cyp735a* contributes to shoot growth both locally, and from the root (both combinations are reduced relative to wild-type self-grafts). We additionally confirmed that *lhk2* and *lhk3* contribute to shoot elongation principally through their action in the shoot (Gifu shoots on *lhk2 lhk3* roots are significantly longer than both *lhk2 lhk3* shoots grafted to gifu roots, and the *lhk2 lhk3* self-graft).

**Figure 7.**
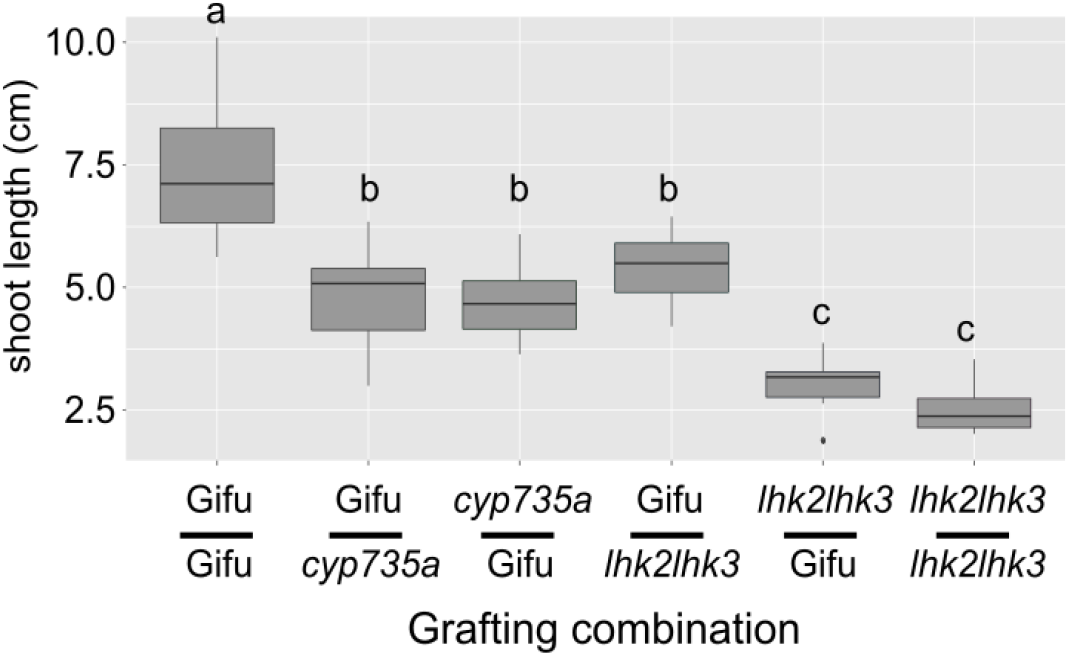
Shoot growth responses in grafted plants. Shoot length was determined for each of the genotype combinations indicated (scions top, roots bottom). Letters indicate significant differences determined using Tukey post-hoc analysis. *n* = 7-13

## Discussion

Cytokinins play critical roles in coordination of root and shoot growth. Here we have shown that cytokinin export from root nodules is required for normal shoot elongation in legumes when relying on symbiotic fixed nitrogen. Mechanistically, cytokinin in the nodule is exported by PUP1, a membrane localised exporter of cytokinins (Fig 8). Shoot localised cytokinin receptors, predominantly *Lhk2* and *Lhk3*, are then required locally for shoot elongation. We identified iP and tZ cytokinin bases as transport substrates for PUP1, which suggests that the export profile reflects the most abundant cytokinin species in nodules. Our grafting experiments with *cyp735a* mutants indicate that tZ bases are produced both locally in the nodule and in the shoot.

**Figure 8.**
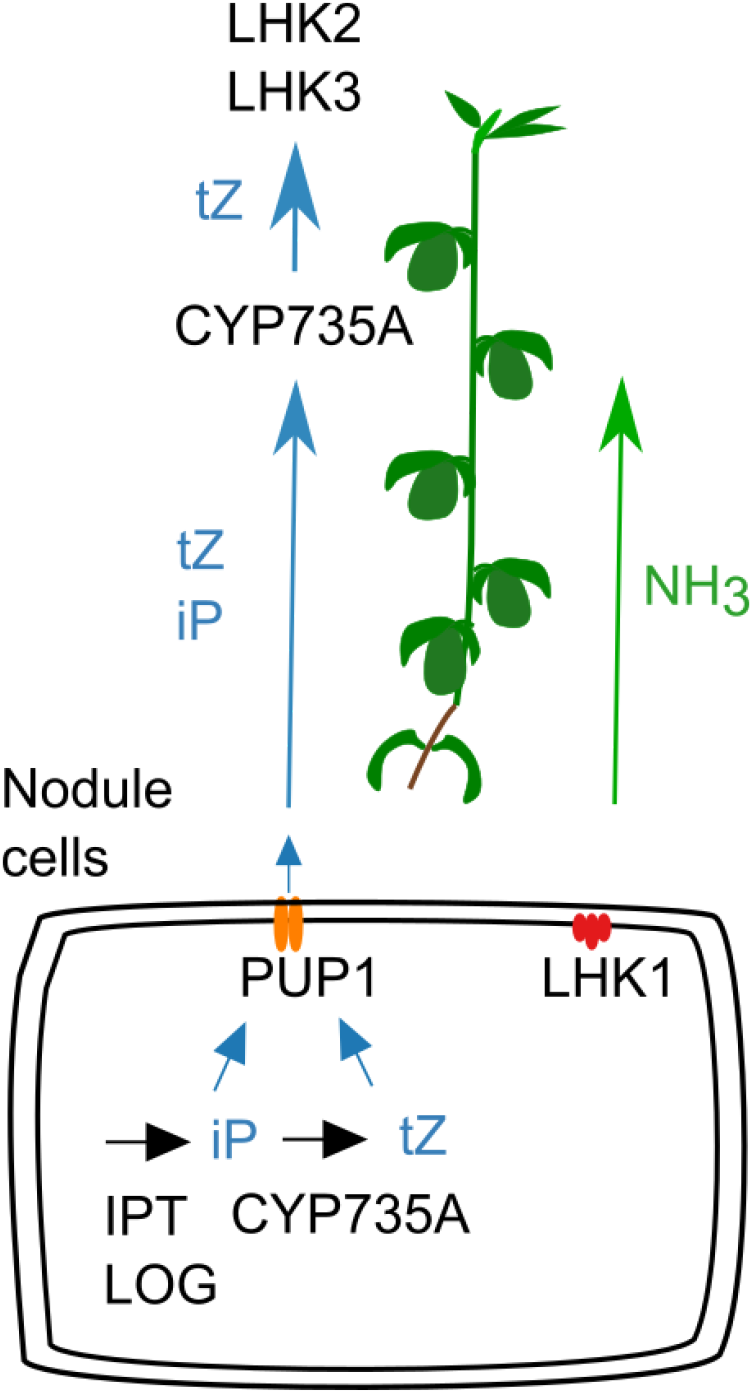
Model of cytokinin export from root nodules. *Pup1* dependent Cytokinin export (both iP and tZ) locally reduces some cytokinin species and signalling outputs. Long-distance cytokinin stimulation of shoot growth depends on the *Lhk2* and *Lhk3* receptors functioning in the shoot and is primarily a tZ dependent effect. Activity of *Cyp735a* in both the nodule and shoot contribute to tZ dependent shoot elongation response.

Stimulation of shoot elongation by PUP1 exported cytokinin is supported by the fact that *pup1* mutants have reduced shoot growth. This reduction occurs despite no difference in nitrogen fixation and equivalent nodule numbers to wild type in at least one *pup1* allele. *PUP1* transcripts are also detected almost exclusively in nodules, with no RNAseq reads in any shoot tissues, and only minor expression in roots outside of nodules. At the same time, expression of the cytokinin hydroxylase *Cyp735a* reaches the highest levels in mature nodules, indicating production of tZ is likely to peak in this tissue. The high transcript abundances for both *Pup1* and *Cyp735a* in mature nodules supports the fact that this tissue is a major cytokinin export organ at this developmental stage. Cytokinin signalling as determined by TCS activity has also begun to decline in the nodule at this point (Held et al., 2014), indicating that the high CYP735A activity is not having major local effects on cytokinin signalling. Our model for long-distance regulation of shoot growth by cytokinin is reminiscent of similar signalling functions in nitrate signalling reported in other species (Rahayu et al., 2005; Krouk, 2016; Landrein et al., 2018). Our findings suggest legumes, which can obtain nitrogen both from the soil and from nitrogen fixation, use cytokinin to coordinate shoot growth with fixed nitrogen. How legumes integrate nitrate with cytokinin in the context of long-distance signalling requires further investigation as we have previously observed a negative relationship between nitrate and cytokinin biosynthesis within the nodule (Lin et al., 2021).

Mutants in *pup1* display intermediate phenotypes relative to the more severely compromised *cyp735a* and *lhk2 lhk3* mutants. This may reflect simple redundancy, where other transporters, such as *Pup4* and *Pup5* which show compensatory expression in *pup1* mutants, contribute to cytokinin export. The transport direction and subcellular localisation of *Pup4* and *Pup5* however remains to be determined. Members of other transporter classes, such as ABCG family transporters, may also support systemic cytokinin transport as is seen in Arabidopsis (Zhang et al., 2014). The broad expression of *Pup1* in all nodule cells indicates that it is not likely to be directly loading cytokinin into xylem tissues for export. One scenario may be that *pup1* exports cytokinins to the apoplast and other vascular localised transporters are required for export to the shoot. This could include PUPs or members of other families of cytokinin transporters. Recently ABCG family transporters have been shown to play important roles in symbiotic cytokinin signalling (Jarzyniak et al., 2021), and organ formation (Jamruszka et al., 2022) and members of this family may be candidates for additional roles later in the symbiosis.

Cytokinin signalling may be activated through cytokinin receptors localised to either the plasma membrane or endoplasmic reticulum (Zürcher et al., 2016; Antoniadi et al., 2020; Kubiasová et al., 2020). Cytokinin response in *pup* mutant nodules appears to have a minor reduction in mutants, as became evident in both RNAseq profiles and TCSn experiments. This suggests that cytokinin export from the cell has a role in stimulating cytokinin signalling in the nodule and indirectly suggests that extracellular cytokinin concentration is important in determining cytokinin signalling within legume nodules. Our demonstration that *LjPup1* functions as a cytokinin exporter contrasts with the demonstrated import function of AtPUP14 (Zürcher et al., 2016). Further structural or biochemical studies will be required to identify the critical features determining directionality of PUP transporters, which will be essential to describe the impacts on cytokinin homeostasis and localisation in different contexts.

In conclusion, our results demonstrate that cytokinin export from symbiotic root nodules plays a role in coordinating shoot growth with the supply of fixed nitrogen. Additionally, we have identified new players in long distance transport of cytokinins, with the demonstration of specific cytokinin export by a member of the PUP family.

## Acknowledgements

This work was supported by grants from the Swiss National Funds (project 31003A_165877 and 310030_197563 to MG. YC was supported by the Chinese Scholarship Council. We are grateful for support from the project Engineering Nitrogen Symbiosis for Africa (ENSA) currently supported through a grant to the University of Cambridge by the Bill & Melinda Gates Foundation and UK government’s Department for International Development (DFID). Analysis by YR and WK is supported by Netherlands Organization for Scientific Research (VENI863.15.010).

## Author contributions

Conducted experiments Yumeng Chen, Jieshun Lin, Flavien Buron, Dugald Reid; Cytokinin measurements Yuda Purwana Roswanjaya, Wouter Kohlen; Microscopy Marcin Nadzieja; Tobacco transport and localization assays Jie Liu, Markus Geisler; Supervision and project conception Jens Stougaard and Dugald Reid; Wrote paper Dugald Reid; Revised paper all authors.

## Data availability

RNAseq data for wild-type and *pup1* mutant nodules is available at NCBI under Bioproject PRJNA622396.

## Supporting information

**Supplemental table 1.**
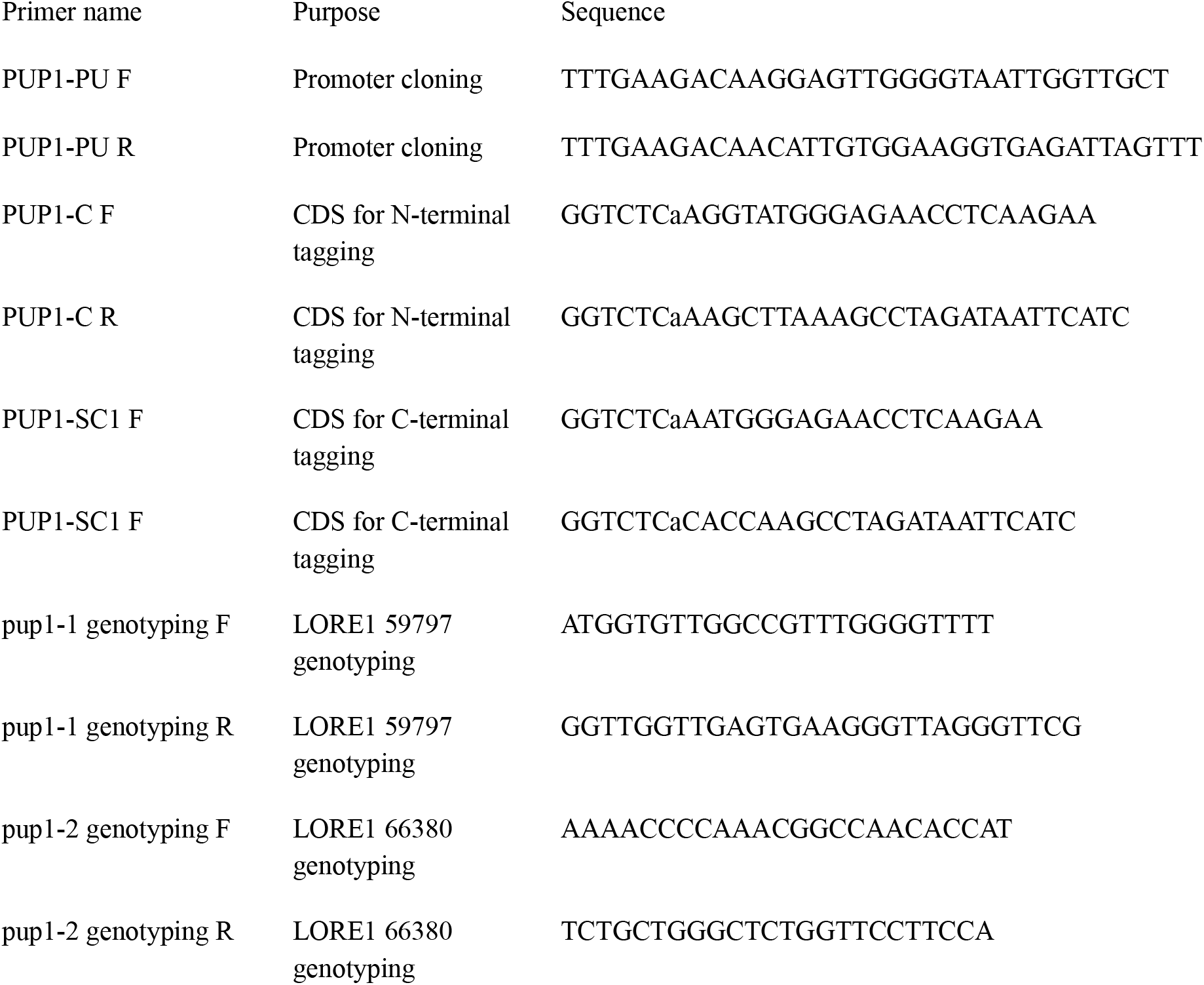
Primers used in this study

**Supplemental dataset 1**. RNAseq normalised reads and differential expression statistics

## Supplemental figures

**Figure S1.**
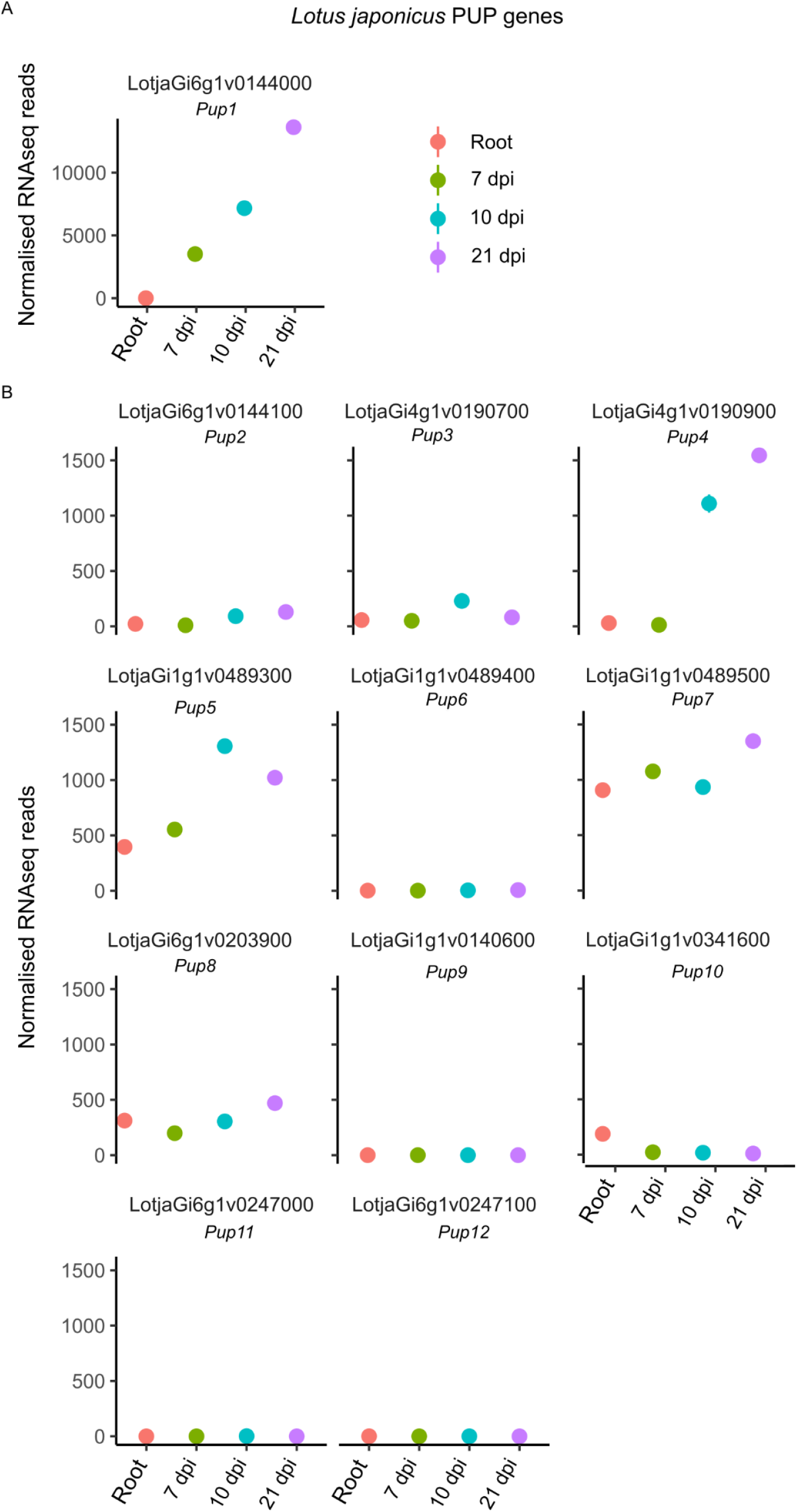
PUP family transcript levels in nodules. Normalised RNAseq transcript levels for *Pup1* (A) and *Pup2-Pup12* (B) obtained from the RNAseq expression atlas at lotusBase (lotus.au.dk). Note a different scale is used in A and B. Circles represent the average normalised transcript levels for 3 replicates per condition (1 replicate at 21 dpi).

**Figure S2.**
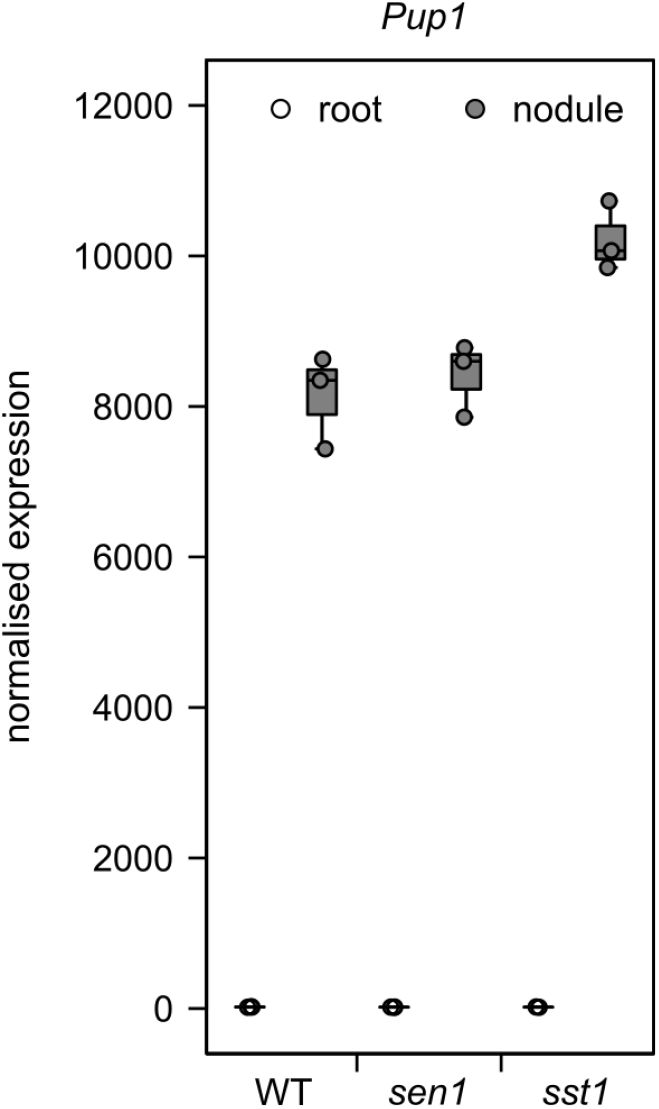
Expression of *Pup1* in *L. japonicus* nitrogen fixation mutants. Data was obtained from microarray based expression analysis of wild-type, *sen1* and *sst1* nodules 21 dpi (Høgslund et al., 2009).

**Figure S3.**
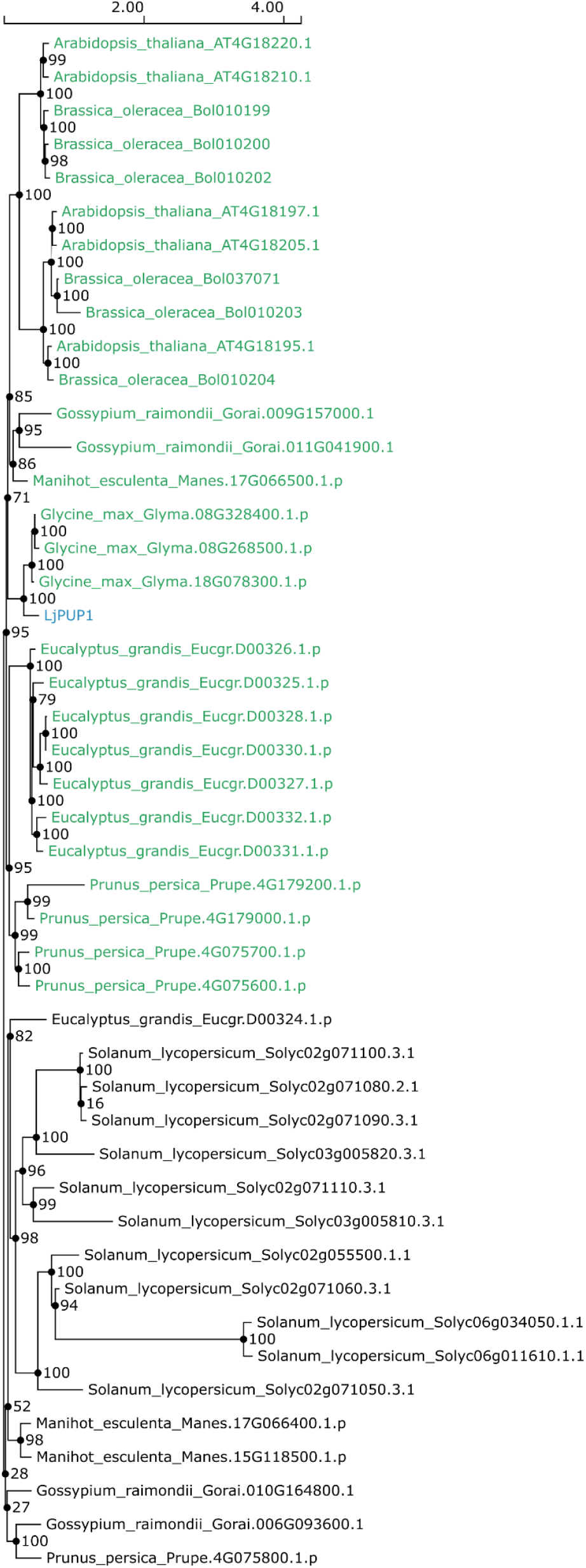
Phylogeny of *Pup1* generated by SHOOT. A PUP1 phylogeny was inferred using the protein sequence of LjPUP1 as SHOOT input (Emms and Kelly, 2022). The orthogroup containing PUP1 is highlighted with green text.

**Figure S4.**
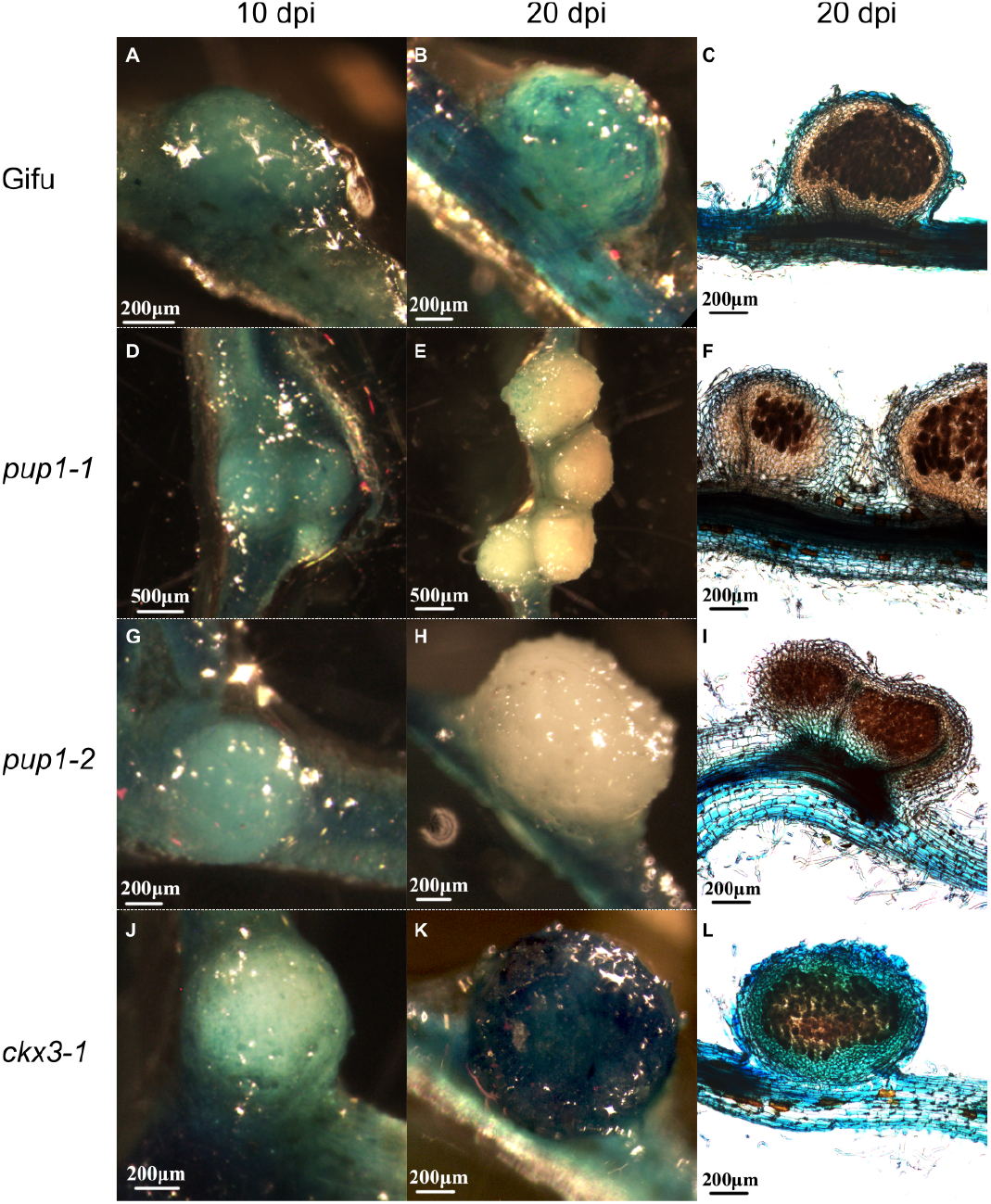
TCSn:GUS in *pup* mutants. TCSn:GUS hairy roots in WT (A-C); *pup1-1* (D-F); *pup1-2* (G-I); *ckx3-1* (J-L). Whole mounts are shown at 10 dpi and 20 dpi and sections from 20 dpi.

**Figure S5.**
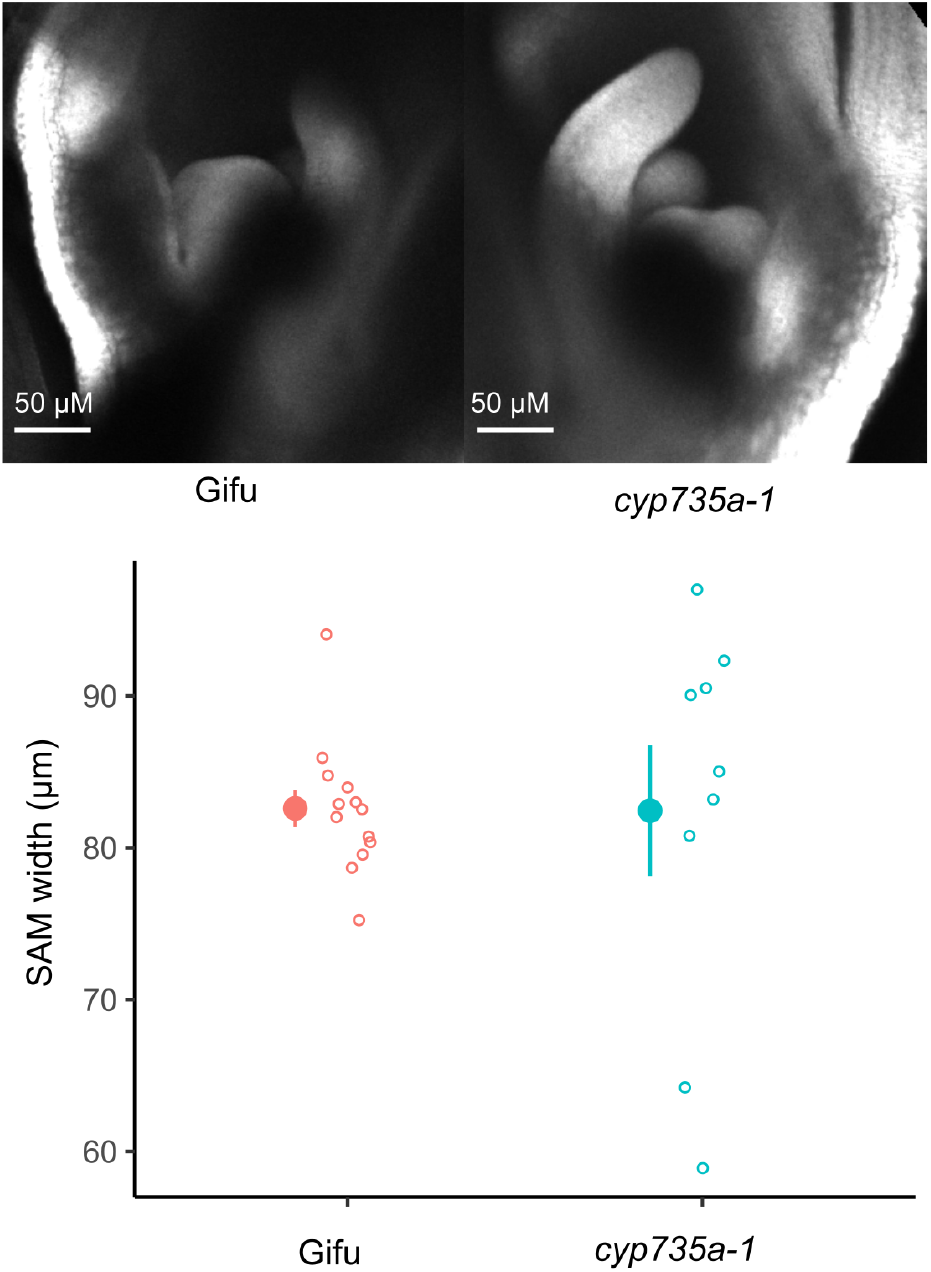
Shoot apical meristem in *cyp735a-1* mutant. Meristems were imaged by confocal microscopy and meristem width measured from the obtained images. *n* = 9-13

